# *De novo* transcriptome assembly, annotation, and identification of low-copy number genes in the flowering plant genus *Silene* (Caryophyllaceae)

**DOI:** 10.1101/290510

**Authors:** Yann J K Bertrand, Anna Petri, Anne-Cathrine Scheen, Mats Töpel, Bengt Oxelman

## Abstract

Phylogenetic methods that rely on information from multiple, unlinked genes have recently been developed for resolving complex situations where evolutionary relationships do not conform to bifurcated trees and are more adequately depicted by networks. Such situations arise when successive interspecific hybridizations in combination with genome duplications have shaped species phylogenies. Several processes such as homoeolog loss and deep coalescence can potentially hamper our ability to recover the historical signal correctly. Consequently the prospect of reconstructing accurate phylogenies lies in the combination of several low-copy nuclear markers that when, used in concert, can provide homoeologs for all the ancestral genomes and help to disentangle gene tree incongruence due to deep coalescence events. Expressed sequence tag (EST) databases represent valuable resource for the identification of genes in organisms with uncharacterized genomes and for development of molecular markers. The genus *Silene* L. is a prime example of a plant group whose evolutionary history involves numerous events of hybridization and polyploidization. As for many groups there is currently a shortage of low-copy nuclear markers, for which phylogenetic usefulness has been demonstrated. Here, we present two EST libraries for two species of *Silene* that belong to large phylogenetic groups not previously investigated with next generation technologies. The assembled and annotated transcriptomes are used for identifying low copy nuclear regions, suitable for sequencing.

## Introduction

*Silene* L. belongs to the carnation family Caryophyllaceae Juss. and comprises about 600 - 700 species (Melzheimer 1980, Zhou et al. 2001, Morton 2005), distributed primarily across the Northern hemisphere. A subdivision of *Silene* into two major clades, subgenus *Silene* and subgenus *Behenantha* (Otth) Endl (=*Behen* (Moench) Bunge) is well supported from a number of molecular studies (Oxelman *et al.* 2001, Popp & Oxelman 2004, Popp & Oxelman 2007, Erixon & Oxelman 2008, Frajman *et al.* 2009, Jenkins & Keller 2010, Rautenberg *et al.* 2010, Petri & Oxelman 2011). The two subgenera contain several hundred species each (Oxelman *et al.* 2011). The majority of the *Silene* species are diploid with x = 12, but allopolyploidy has been found to be common for example within section *Physolychnis* (Benth.) Bocquet (subgenus *Behenantha*), in which the majority of the species are polyploid (Popp et al. 2005; Popp & Oxelman 2007; Petri & Oxelman 2011).

In addition to the phylogenetic studies, *Silene* receives considerable interest as a model in evolutionary and ecological research (e.g., Bernasconi *et al.* 2009). Although a completely sequenced genome has yet to be published, several genome-wide data sets are becoming available through the advent of Next Generation Sequencing technologies. Blavet *et al.* (2011) generated transcriptome data from 1 - 3 individuals of *S. latifolia* Poir., *S. dioica* (L.) Clairv., *S. vulgaris, S. marizii* Samp., and *Dianthus superbus* L., (publically available at *http://www.siesta.ethz.ch*). Sloan *et al.* (2012) published transcriptome data from one specimen of each of three geographically separate populations of *S. vulgaris* (Moench) Garcke (*http://silenegenomics.biology.virginia.edu*). Transcriptomic data from *S. latifolia* has also been generated by Bergero *et al.* (2011) and Muyle *et al.* (2012). All four investigated species of *Silene* belong to subgenus *Behenantha*. *Silene vulgaris* belongs to section *Behenantha* within subgenus *Behenantha* (Oxelman *et al.* 2011), whereas the three other *Silene* species belong to section *Melandrium* in the same subgenus (Rautenberg *et al.* 2010, Oxelman *et al.* 2011).

Low-copy nuclear genes have proved suitable for phylogenetic reconstruction in difficult groups where successive interspecific hybridizations in combination with genome duplications (polyploidy) have produced reticulate interspecific relationships (e.g. Popp & Oxelman 2004, Popp *et al.* 2005, Petri & Oxelman 2011, Marcussen *et al.* 2012). Although key for disentangling these complex phylogenies, there is a shortage of low-copy nuclear markers, for which phylogenetic usefulness has been demonstrated. The limitation is partly technical as the presence of more than one allele and similar loci render the Sanger sequencing step problematic. Therefore, it is desirable to identify conserved regions of low-copy number genes where PCR primers or probes for sequence capture can be designed. In order to increase the resources of low-copy nuclear markers available for phylogenetic studies across *Silene*, we have amassed 454 transcriptome data from two diploid species located in different subgenera: *S. uralensis* (Rupr.) Bocquet, (section *Physolychnis*, subgenus *Behanantha*), and *S. schafta* J.G.Gmel. ex Hohen. (section *Auriculatae* (Boiss.) Schischkin, subgenus *Silene*). We compared the acquired transcriptome sequences in order to identify cross species genetic markers candidates. A number of primers for low copy genes were designed and tested across *Silene* species for their phylogenetic utility. The assemblies are processed by annotating the sequences with gene names and Gene Ontology terms and by identifying unique transcripts that were not captured by previously published transcriptomes..

## Material and methods

### Library preparation

RNA was extracted from one individual of *S. uralensis* (Rupr.) Bocquet subsp. *arctica* (Th. Fr.) Bocquet, grown from seeds collected in Svalbard, Endalen, 52 m above sea level, and cultivated in the phytotrone at Tøyen, University of Oslo (Gustavsson 14, LG09-S-14-01 to LG09-S-14-10, vouchers at herbarium O), and one of *S. schafta* J.G.Gmel. ex Hohen. from the Botanical Garden in Göteborg (voucher: Oxelman 2565, deposited at herbarium GB), using the mirVana miRNA isolation kit (Ambion) by vertis Biotechnologie AG (http://www.vertis-biotech.com). As many stages of the life-cycle as possible were used (i.e., roots, stem, old and young leaf buds, flowers, developing fruits), ensuring as complete mRNA coverage as possible. The resulting EST libraries were normalized by one cycle of denaturation and reassociation of the cDNA, resulting in N1-cDNA. Reassociated ds-cDNA was separated from the remaining ss-cDNA (normalized cDNA) by passing the mixture over a hydroxylapatite column. After hydroxylapatite chromatography, the ss-cDNA was amplified with 11 PCR cycles. The cDNAs in the size range of 500–700 bp were eluted from preparative agarose gels, tagged by species -specific barcodes, and sequenced on a half picotiter plate on a 454 GS-FLX sequencer with Titanium reagents (Roche) at the Norwegian Sequencing Center (http://www.sequencing.uio.no).

### EST processing

Sequences from the Roche 454 instrument output in Standard Flowgram Format (sff) were converted into fastq format. Adapters were removed using cutadap (Martin 2011), a trimming tool that is tolerant for insertions and deletions within homopolymers. Reads were further filtered to minimum quality (phred score 20) and minimum length (100bp) using the Filter FASTQ (v. 1.0.0) utility from Galaxy (Giardine *et al.* 2005, Blankenberg *et al.* 2010, Goecks *et al.* 2010).

Newbler v. 2.5 (Roche) was used for transcriptome assembly. GS De Novo assembler was run with the “-cdna” option to assemble transcriptomes and the following settings: minimum overlap length = 40, minimum overlap identity = 90, alignment identity score = 2, and alignment difference score = -3. Both isotigs and contigs were generated. The former attempts to represent the original sample’s mRNA and contain splice variants that entered the transcriptome annotation step, while the latter aim at capturing exons that were used for finding potentially useful regions for designing primers or probes of low copy number genes. In addition, Newbler groups related isotigs into isogroups that potentially represent all transcripts derived from a single gene.

EST reads that did not build contigs were added to the isotigs. Sequences derived from chloroplast and mitochondria were filtered against a custom database containing all fully sequenced chloroplasts and mitochondria for Viridiplantae using BLAST (Altschul *et al.* 1990) similarity search (BLASTN program with e-value < 1e-6).

### Candidate genes identification and primers design

Following sequence assembly, all contigs were analyzed using GeMprospector (Fredslund *et al.* 2006; http://cgi-www.daimi.au.dk/cgi-chili/GeMprospector/main), where the first step is to align the input EST data to the existing ESTs in a database consisting of genomic sequences from *Arabidopsis thaliana, Lotus japonica*, and *Medicago truncatula*, and then use BLASTX to identify *Arabidopsis* orthologs. ESTs with only one or two hits in the *Arabidopsis* proteomewere selected as potential candidates. Intron-exon boundaries were tagged based on comparisons with genomic sequences from *Lotus japonicus* and *Medicago trunculata*. The final step in the pipeline is defined primers located at suitable distances of either side of the putative intron-exon boundaries. This was done using the PriFi software (Fredslund *et al.* 2005), which suggests cross-species conserved primer sites.

### Primer optimization

In order do test the usefulness of some of the primers suggested by GeMprospector, all first ranked primers were blasted against local databases of the two transcriptomes using TBLAST and TBLASTX in blast v.2.2.23+ (Altschul *et al.* 1997). Primers that gave one (and only one) significant BLAST hit were made degenerate to match both *S. uralensis* and *S. schafta*.

These primers were optimized for their PCR amplification efficiency, covering the six major subgroups of the genus *Silene*. DNA was extracted for two specimens of each subgroup; *S. uralensis* (subgenus *Behenantha* section *Physolychnis*), *S. schafta* (subgenus *Silene* section *Auriculatae*), *S. latifolia* Poir. (subgenus *Behenantha* section *Melandrium), S. cryptoneura* Stapf, (subgenus *Behenantha) S. nutans* L. (subgenus *Silene* section *Siphonomorpha), S. aegyptiaca* L.f (subgenus *Behenantha* section *Atocion)* as described in Petri & Oxelman (2011). See table X for voucher information. PCR amplifications were performedusing PHUSION polymerase (Finnzymes) using the manufacturer’s instructions. All reactions were run with an annealing temperature gradient ranging from 59°C to 71°C. (Reaction results and PCR programme details are avaliable on http://www.sileneae.info/Boxtax) Primer pairs producing single bands on a 1.5% agarose gel from at least four of the major *Silene* groups were selected for further processing.

These were used for amplification of all the above listed specimens in a second PCR, using the lowest annealing temperature which produced a single sharp band during the first amplification. The products were sequenced with Sanger sequencing reactions performed by Macrogen Inc. (www.macrogen.com).

### Isotigs and singletons annotation

A similarity search against non-redundant protein reference sequence database (Pruitt *et al.* 2005) RefSeq Release 54 was performed with BLASTX and only isotigs and singletons with a significant similarity score (e-value < 1e-5) entered the annotation procedure. Gene ontology (GO) annotation was carried out with Blast2go (Conesa & Gotz 2008). During the initial mapping, candidate GO terms were retrieved from GO terms associated with the 5 first blast hits (e-value < 1e-6). Sequences were annotated with the default parameters and augmented with the ANNEX function. The resulting annotation was probed for biological functions unambiguously associated with non-plant organisms. These sequences were filtered out after manual inspection.

### Similarity search against published Silene and Dianthus species

We built a custom BLAST database (Silene_db) by concatenating the publicly available *Silene* and *Dianthus* transcriptome data from Bergero *et al.* (2011), Blavet *et al.* (2011) and Sloan *et al.* (2012). After removing short entries (< 100bp), the resulting set comprises 434576 sequences distributed accross five species. Searching against this database (BLASTN, e-value < 1e-9) allowed the identification of expressed genes that were not previously obtained in transcriptome studies within *Silene* and *Dianthus*.

*Reciprocal BLAST comparisons between* S. uralensis, S. schafta *and* S. vulgaris Transcriptomes comparisons were performed on contigs instead of isotigs. For each species, contigs were clustered and their consensus sequences computed with UCLUST (Edgar, 2010). A minimum identity value of 95% was used for clustering after preliminary tests showed that the value clustered accurately transcripts in groups containing alleles and sequences with homopolymer errors while separating paralogs. The consensus sequences were then subjected to a Reciprocal BLAST analysis wherby all best matches between the three trancriptome datasets from *S. uralensis, S. schafta* and *S. vulgaris* were identified using the program reciprocal_blast-nt.py available at https://github.com/mtop/reciprocal_blast. During each pairwise comparison of transcriptome sequences, we assigned best matching pairs based solely on highest e-values.

## Results

The 454 run resulted in 319 341 reads (average read length 376.1±129.5) for *Silene uralensis*, and 312084 reads (average read length 375.6±129.3) for *Silene schafta*. After quality and adapter trimming 78.38% and 77.85% of reads remain for *S. uralensis* and *S. schafta* respectively. Their assemblies produced 30748 contigs (average read depth of 5.6) for *S. uralensis*, and 27740 contigs (average read depth of 5.9) for *S. schafta* (Table 1). All contigs and primer site suggestions from GeMprospector are available at http://www.sileneae.info/Sileneae_project/GEM_EST.html. In total, 70 primer sites were suggested from the *S. schafta* transcriptome, and 88 from the *S. uralensis* transcriptome. 34 primer pairs were chosen to be tested for initial amplification. 20 primer pairs that generated single bands across the at least four of the subgroups were used for a second amplification and sequencing. In general, the primer sequences proposed by GemProspector need to be refined with respect to primer length, codon degeneracy and position, and specificity to the taxa addressed.

The proportion of isogroups that contain putative splice variant of a gene, i.e. isogroups that possess more than one isotig, was 10% for *S. uralensis* and 9% for *S. schafta* (Table 2). The gene space covered by our transcriptomes was accessed through a GO annotation procedure that characterized 26% of isotigs and singletons for *S. uralensis* and 28% for *S. schafta* (Table 3).

A large proportion of assembled sequences lack significantly similar counterparts in the *Silene* and *Dianthus* sequences database. Among these new gene transcripts for Caryophyllaceae, 1109 sequences for *S. uralensis* and 4386 for *S. schafta* were functionally annotated. For *S. uralensis* these transcripts relate principally to metabolic (50%), mainly nitrogen metabolism, and cellular processes (46%), whereas for *S. schafta* unique transcripts are subdivided into metabolic (48%), mainly primary metabolic processes (34%) and cellular processes (41%). Only a handful of new gene transcripts (8 for each species) were likely to be specifically expressed in non-floral organs.

The *Silene uralensis* and *S. schafta* transcriptomes were compared to the published data from *S. vulgaris*, which is the most comparable dataset in term of sequencing technology, assembly and coverage. Initial comparisons performed on isotigs and singletons yielded very low similarity scores between the three transcriptomes (see http://matstopel.se/notebook/Reciprocal_BLAST_S.schafta-S.uralensis-S.vulgaris for more details). This result could be explained by the presence of factors susceptible to confound a reciprocal BLAST analysis such as alternative splicing, sequencing errors and allelic variation. We circumvented the alternative splicing problem by performing transcriptome comparisons on consensus sequences from contigs that had been clustered according to sequence similarity, which in ideal conditions should capture exon sequences, instead of using isotigs that represent splice alternative sequences. The other factors were handled by grouping sequences in clusters of similar sequences and blasting their consensus sequences. Selecting the appropriate level of similarity that will cluster alleles and contigs with sequencing errors while keeping paralogs separated is a critical step. Admittedly, a single similarity value would not perform equally well in different species and on genes with different histories and functional constrains. Nevertheless, visual inspection of a sample of clusters in each species indicated that a 95% similarity value did not cluster contigs that patently corresponded to paralogs and that group members were in majority sequencing error variants. Contigs clustering using UCLUST significantly reduced the number of sequences in each set (Table 4), albeit keeping a large proportion of contigs as singletons and hence demonstrating that Newbler produced efficient assemblies despite the presence of sequencing errors and distinct alleles. The reciprocal BLAST analysis revealed that a high number of consensus sequences (60%, 63% and 80% from *S. vulgaris, S. schafta* and *S. uralensis* respectively) had a single best match in all three transcriptomes (Fig. 1). Results from the individual analyses are available at http://dx.doi.org/10.6084/m9.figshare.96750.

**Fig. 1.**
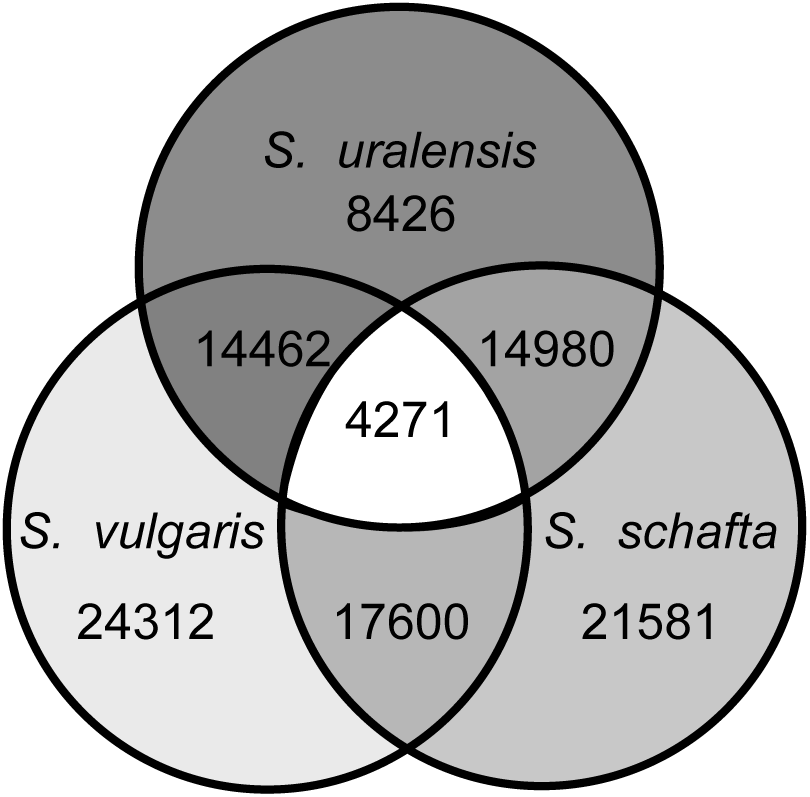
Results from the reciprocal BLAST analysis of consensus sequences from clustered contig sequences from the three species *Silene schafta, S. uralensis* and *S. vulgaris*. The Venn-diagram shows the number of reciprocal best matches between pairs of datasets, as well as the number of best matches between all three datasets (in the center of the figure).

The annotated sequences can be accessed at http://www.sileneae.info/blast/db/

## Discussion

Our data includes a large proportion of transcripts with a significant similarity to other *Silene* and *Dianthus* sequences that represent putative orthologs which could be used after further orthology analyses for phylogenetic inference. Furthermore, a significant amount of transcripts was not captured by previous studies. As singletons dominate among these sequences, their lack of similarity with other *Silene* and *Dianthus* transcripts could be due to pyrosequencing errors or they might represent repeated elements that are abundant in *Silene* genomes (Macas *et al.* 2011). An alternative explanation could emphasize the differences in the cDNA library preparations. We have followed a similar protocol to Sloan *et al.* (2012) for collecting tissues from leaf, root and floral parts and normalized our libraries, whereas the transcriptomes generated by Blavet *et al.* (2011) are derived from solely floral buds without library normalization. Normalization reduces redundancy in the cDNA pool thereby resulting in more even coverage and allowing rare transcript to be sequenced. It is likely that our unique transcripts refer to genes expressed in other plant tissues than floral buds at the time of the plant material harvesting or that were present at low expression levels.

There is an ongoing revolution in the field of molecular phylogenetics as vast amounts of data are amassed through next generation sequencing (NGS) methods such as transcriptome acquisition (e.g. Roeding *et al.* 2009) and restriction-site associated DNA (RAD) sequencing (e.g. Rubin *et al.* 2012) while phylogenomic reconstructions based on full genome comparisons are likely to be performed in a foreseeable future. Reciprocal BLAST analyses confirmed that our transcriptomes generated high levels of sequence similarity with *S. vulgaris*. A total of 4271 consensus sequences from clustered contigs had a best match in all three datasets, and hence represent putative orthologs that need to be validated with phylogenetic methods.

For the time being however, these mass sequencing techniques are still not routinely applied in investigation encompassing large sets of taxa and in studies without access to well preserved DNA such as extractions from herbarium specimens. Consequently many evolutionary studies are still relying on a set of few selected genes, which can nevertheless be sequences with NGS once specific PCR primers or probes for sequence capture have been designed.

Together with the results from other recent comprehensive *Silene* transcriptome studies (Blavet *et al.* 2011, Sloan *et al.* 2012), our data contributes to the resources available for research in *Silene*. In particular, our data closes large taxonomic gaps, by including one representative from the large subgenus *Silene* clade (*S. schafta*), as well as a representative from the species-rich *Physolychnis* clade (*S. uralensis*). Although there is much interest in *S. vulgaris* and *S. latifolia,* which both belong to subgenus *Behenantha*, they merely represent two of many lineages stemming from an ancient radiation (Erixon & Oxelman 2008). Many of the general evolutionary questions raised in *Silene* requires a firm understanding of the phylogenomic relationships in the genus and the currently available data provide a good backbone for further transcriptome studies of other *Silene* species, once they are obtained.

## Acknowledgements

The authors wish to thank Lovisa Gustavsson for sharing parts of her Svalbard collection. Financial funding was obtained from grants to AP from Helge Ax:son Johnssons fund and to BO from the Swedish Research Council.

